# Elucidating biosynthetic pathway of piperine using comparative transcriptome analysis of leaves, root and spike in *Piper longum* L

**DOI:** 10.1101/2021.01.03.425108

**Authors:** Prem Kumar Dantu, Mrinalini Prasad, Rajiv Ranjan

## Abstract

*Piper longum* (Pipli; Piperaceae) is an important spice valued for its pungent alkaloids, especially piperine. Albeit, its importance, the mechanism of piperine biosynthesis is still poorly understood. The Next Generation Sequencing (NGS) for *P. longum* leaves, root and spikes was performed using Illumina platform, which generated 16901456, 54993496 and 22900035, respectively of high quality reads. In *de novo* assembly *P. longum* 173381 numbers of transcripts were analyzed. Analysis of transcriptome data from leaf, root and spike showed gene families that were involved in the biosynthetic pathway of piperine and other secondary metabolites. To validate differential expression of the identified genes, 27 genes were randomly selected to confirm the expression level by quantitative real time PCR (qRT-PCR) based on the up regulation and down regulation of differentially expressed genes obtained through comparative transcriptome analysis of leaves and spike of *P. longum*. With the help of UniProt database the function of all characterized genes was generated.

## Introduction

*Piper longum* popularly known as pippali (family: Piperaceae) is an important medicinal plant ranking third after *P. nigrum* (black papper) and *P. betle* (betel) in economic importance (Jose and Sharma, 1985). It grows extensively in hot and humid Indo-Malaya region (Manoj *et al*., 2004). *P. longum* has several important bioactive compounds viz. alkaloid, flavonoids, glycosides, tannins, phenols and sterols (Rami *et al*., 2013). Piper fruit has several alkaloids, piperine being the most important one and is the reason for its pungency (Zaveri *et al*., 2010). The plant is a powerful stimulant for both digestive and respiratory systems and has a rejuvenating effect on the lungs (Charaka by Sharma, 1996). *Piper* fruit (spikes) plays an important role in multiple aliments as increased thyroid hormone level, thermogenic response, immune stimulatory, anti-ulcer, anti-amoebic, anti-oxidant, hepatoprotective, cardiovascular and anti-inflammatory activities (Dahanukar and Karandikar, 1984; Warrier *et al*., 1995; Chauhan *et al*., 2011; Sharma *et al*., 2012; Prasad *et al*., 2018). The abundance of medicinal value of this plant resulted in extensive and indiscriminate collection from the wild threating the very existence of it necessitating developing methods to conserve *P. longum* (Nair 2000; Rani and Dantu, 2012).

Collection pressure on *P. longum* could be relieved by either finding alternate source plants or through biotechnological intervention where piperine content in leaves could be increased. Piperine accumulation has been found to be highest in the spikes, followed by roots and least amount has been detected in the leaves of *P. nigrum* (Semler *et al*., 1988) and *P. longum* (Prasad *et al*., 2018, 2019). However, piperine biosynthesis in *P. longum*, its transportation within the plant and storage are still poorly understood. Although, Piperidine conversion to piperine using Piperoyl-CoA has been worked out in *P. nigrum* and steps of lysine conversion to piperidine have been elucidated in the bacterium *Pseudomonas aeruginosa* (Fothergill and Guest, 1977; Greisler and Gross *et al*., 1990; Bunsupa *et al*., 2012).

Despite the fact that the acetyl or malonyl CoA-based reaction of feruloyl-CoA to be a conceivable component for the C2-extention of the cinnamic acid-derived precursor ensuring piperoyl-CoA development. In these steps no data is available regarding molecular and enzymatic aspect. In another case, the related amide capsaicin derived from buturyl-CoA by C2-elongations which are encoded by Pun1 gene (Stewart et al., 2005). Explanation of detailed information of genome related to the pungency of capsaicin (Kim et al., 2014).

Recent advances in transcriptome analysis using Next Generation Sequencing has elucidated metabolic pathways in several medicinal plants, such as, *Artemisia annua* (Paddon and Keasling, 2014), *Withania somnifera* (Gupta *et al*., 2015), *Panax ginseng* (Zhang *et al*., 2017), *Picrorhiza kurroa* (Shitiz *et al*., 2015) to name a few. Transcriptome analysis of the seed (Hu *et al*., 2015) and root (Gordo *et al*., 2012) of *P. nigrum* has generated a large data bank of gene sets involved in various processes. However, such a transcriptomic study in *P. longum* is still lacking. In non-model plants such as, *P. longum*, deep genome sequencing using Next Generation Sequencing is the only viable method for generating large molecular data set that can identify proteins (Yu *et al*., 2011; Low and Heck, 2016; Low *et al*., 2019).

A detailed investigation of the genome of long pepper transcriptome using RNA-seq can give us a great insight into this species. The genomic data generated could be useful in understanding the genes involved in biosynthetic pathway of not only piperine but also other secondary metabolites of importance. The knowledge thus generated could be useful to understand ecology, evolution of this species and also in biotechnological interventions (Ekblom and Galindo, 2011; Bohan *et al*., 2017; Derocles *et al*., 2018). In the present study transcriptome of long pepper leaves, roots and spikes were analyzed using Illumina HiSeq 2000 platform. Approximately 17, 55 and 23 million clean reads were generated from leaves, roots and spikes, respectively resulting in 1,73,381 unigenes. Analysis of the transcriptome data showed gene families that were involved in the biosynthesis of piperine and other secondary metabolites and in housekeeping genes. This is the first dataset of sequence analysis of long pepper leaves, roots and spikes and will be a useful genomic library for any molecular research.

## Material and Methods

### Plant materials

*Piper longum* plants were grown in net house of Department of Botany, Dayalbagh Educational Institute, Agra (U.P.), India. Young leaves, roots and spikes were collected in November from 4 years-old plants with the help of sterilized blade and transferred into RNAlater (Sigma-Aldrich, USA) for future use.

### RNA isolation and library construction

Total RNA was isolated from three different tissues i.e. leaf, root and spike using Trizol (Invitrogen) method and Xcelgen plant RNA kit. The quality of the isolated RNA was checked on 1% Formaldehyde Denaturing Agarose gel and quantified using Nanodrop8000 spectrophotometer. The paired-end sequencing libraries were prepared using Illumina TruSeq Stranded mRNA Library Preparation Kit as per the protocol. Total isolated RNA was subjected to Oligo dT beads to enrich mRNA fragments, then subjected to purification, fragmentation and priming for cDNA synthesis. The fragmented mRNA was converted into first-strand cDNA, followed by second-strand cDNA synthesis, A-tailing, adapter-index ligation and finally amplified by recommended number of PCR cycles. After obtaining the Qubit concentration for the library and mean peak size from Bio-analyzer 2100 (Agilent Technologies), library was loaded into Illumina platform for cluster generation and sequencing. Library quality and quantity check was performed using Agilent DNA High Sensitivity Assay Kit.

### Transcriptome sequencing and analysis

RNA was pooled for sequencing and preparation was multiplexed on a single flow cell of an Illumina TruSeq. The final library was denatured and appropriate working concentration was loaded on to flowcell. Briefly, double stranded cDNA was subjected to Covaris shearing followed by end-repair fragments of overhangs. The fragments were A-tailed, adapter ligated and then enriched by limited number of PCR cycles. Prepared library was loaded into Illumina platform for cluster generation and sequencing was performed using paired-end (PE) 2×150 bp library on Illunima platform (Illumina TruSeq RNA 500) (Bolger *et al*., 2014). Subsequently, all assembled transcript contigs were validated using CLS Genomics Workbench 6 by plotting high quality reads back to the assembled transcript contigs. All CDS (protein coding sequences) were predicted from the assembled transcripts/ scaffolds using Transdecoder (rel16JAN2014) with default parameters.

### Functional annotation and gene ontology (GO) analysis

The functional annotation were performed on the predicted CDS of plant sample by aligning the CDS to non redundant protein database of NCBI using Basic Local Alignment Search Tool (BLASTx) (Altschul *et al*., 1990) with a minimum E-value less than 1e-5. CDSs were predicted with the help of Transdecoder software. BLASTx 2.2.30+ software was used for functional annotation detection standalone. The total CDS was found to have BLASTx hit against ‘NR database of NCBI’. A BLAST explore was performed against the non-redundant protein (NR) and nucleotide sequences (NT) databases in the National Center for Biotechnology Information (NCBI) with an E value less than 1e-5. For annotating the transcript tags, the best match output from each BLAST was used. After each BLAST search annotation tags with no matches and ones with predicted annotations were extracted for next sequential BLAST search. GO assignment was used to classify the functions of the predicted CDS. The GO mapping also provided ontology of defined terms representing gene product properties which were group into three main chategories: molecular function (MF), biological process (BP) and cellular component (CC) (Ashburner *et al*., 2000). GO mapping was carried out using Blast2GO Pro software (Gotz *et al*., 2008) to retrieve GO terms for all the BLASTx functionally annotated CDS which included use of BLASTx result accession IDs to retrieve gene names or symbols.

### Classification of assembled transcriptomes

KAAS (KEGG Automatic Annotation Server - http://www.genome.jp/kegg/ko.html) was used to functionally annotate the CDS by BLAST comparisons against KEGG gene database (Moriya *et al*., 2007). The BBH (Bi-directional best hit) option was used to assign KEGG Orthology (KO) terms. The KEGG Orthology database was used for pathway mapping. The unigenes were assignments of polypeptides produced from combined assembly and mapped into metabolic pathways according to KEGG (Kyoto Encyclopedia of Genes and Genomes).

### Differential gene expression analysis

Differential gene expression analysis has been carried out for a total predicted CDS among (a) *P. longum* leaf (as Control) vs spike (as Tested), (b) *P. longum* leaf (as Control) vs root (as Tested) and (c) *P. longum* root (as Control) vs spike (as Tested) using DESeq package (Wang *et al*., 2010). Criteria which was taken for up regulated filtration is log2FC>2 AND p<0.05 and for down regulated filtration is log2FC<-2 AND p<0.05. Differentially expressed genes identified in *P. longum* leaf, root and spike were analyzed by hierarchical clustering (Eisen *et al*., 1998). A heat map was constructed using the log-transformed and normalized value of genes based on both Pearsonun centered correlation distance and on complete linkage method.

### Transcription factors

For the identification of transcription factor families represented in transcriptome, all the predicted CDS were searched against all the transcription factor protein sequences in the Plant Transcription Factor database (PlnTFDB; http://planttfdb.cbi.pku.edu.cn/download.php) using BLASTx with e-value less than 1e-5.

### Identification of Simple Sequence Repeats (SSRs)

For identification of SSRs, all the assembled transcripts / scaffolds were searched with Perl Script MISA. SSRs generated from transcriptome sequence of leaves, root and spikes of same plants were used in Microsatellite program (MISA). Scrutiny contains microsatellites from di-nucleotide to hexa-nucleotide. Mono-nucleotide repeat motifs were not considered in the analysis due to chances of homopolymer tailing in ESTs generated. Unigenes were used as reference data in SSRs detection. SSRs having a flanking of 150 bp were filtered and used for primer designing (Temnykh *et al*., 2000).

### Quantitative gene expression analysis

The total RNA was isolated from samples using PureLink^®^ RNA Mini Kit (Invitrogen, Thermo Fisher Scientific) and quality of RNA was checked on 1% denatured agarose gel using MOPS buffer. The cDNA synthesis from each RNA samples was carried out by using cDNA synthesis kit SuperScript^®^ III First-Strand Synthesis System for RT-PCR (Invitrogen, US). PCR reactions were set up in 10µl reaction mixture in 96-well PCR plates (Don *et al*., 1991). PCR products were separated on 4% agarose gels (Smith *et al*., 2000). Relative and quantitative expression in leaves and spikes was analyzed using Real-Time PCR Detection Machine (Stratagene Mx3005P, Agilent Technologies) and KAPA SYBR^®^ FAST qPCR Master Mix (2X) kit (KAPA BIOSYSTEMS,US). For relative analysis quantitative PCR was performed using cDNA templates; 10µl of 2X KAPA SYBR FAST qPCR Master Mix; 200nM of each primer (forward and reverse); 1X 50X ROX high/low and rest of nuclease free water for 20µl complete reaction volume. For relative analysis quantitative PCR was performed using the corresponding cDNA templates and SYBR master mix as per kit’s instruction. For each set of primer, a control reaction was also included having no template. GAPDH gene was used as internal control to estimate the relative transcript level of the genes studied (Zhang *et al*., 2018). All the 27 genes including internal gene (GAPDH) was analyzed by qRT-PCR with the help of cycle parameters 95°C for 10 min followed by 40 cycles of 95°C for 30 sec, 55°C for 30 sec, 72°C for 30 sec and for melting curve 95°C for 1 min, 55°C for 30 sec and 95°C for 30 sec. Data from qRT-PCR was analyzed using comparative ΔΔct and fold changes in the transcript level were calculated using 2^-ΔΔct^ method (Livak and Schmittgen 2001 and Ranjan *et al*., 2012) considering the Ct value of GAPDH as the internal control. Each experiment was repeated using three biological replicates and three technical replicates, the data was analyzed statistically (± Standard Deviation).

## Results

### Sequencing and de novo transcriptome assembly

All the plant samples were collected and immediately stored in RNAlater. Total RNA was isolated using Trizol (Invitogen) method. RNAs of equal quality were mixed for Illumina TruSeq RNA sample preparation Kit sequencing. Raw reads of totaling 5,07,04,36,800 from leaves, 8,16,08,80,210 from roots and 6,87,00,10,500 from spikes were obtained. The NGS for *P. longum* leaves, root and spikes were performed using 2×150 bp on the Illumina Platform that resulted in generation of high quality data. The high quality reads statistics of *P. longum* leaves contained 1,69,01,456 paired reads, roots contained 5,49,93,496 paired reads and spikes contained 2,29,00,035 paired reads. *De novo* master assembly of pooled high quality paired end reads of *P. longum* roots, leaves and spikes samples were accomplished and total number of 1,73,381 transcripts were obtained (Table 1). Total transcriptome, N50 and maximum length of transcripts size in the libraries were 99612836, 722 and 12994 bases, respectively. The size distribution of the raw reads and assembled transcripts from libraries were characterized (Figure 1).

**Table 1:**
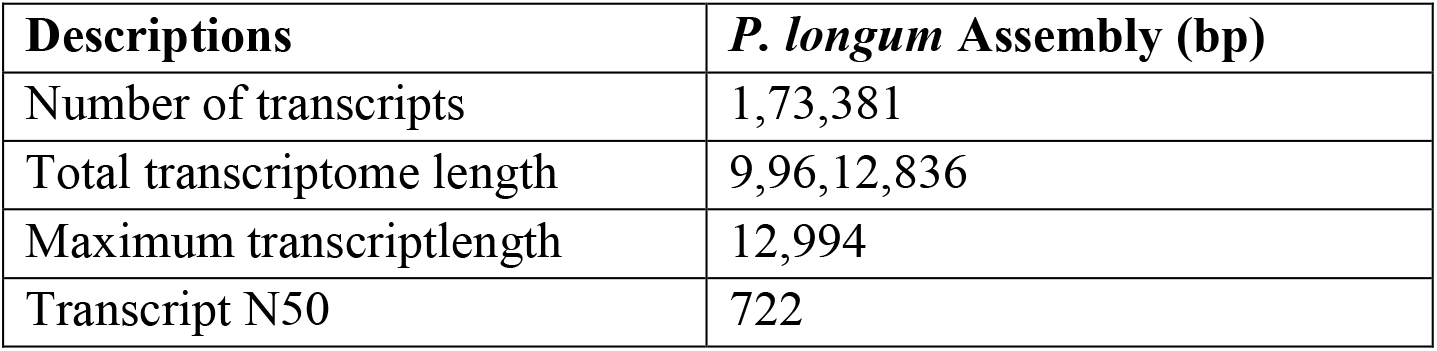
Statistics of assembled transcript of *P. longum* sample.

**Figure 1:**
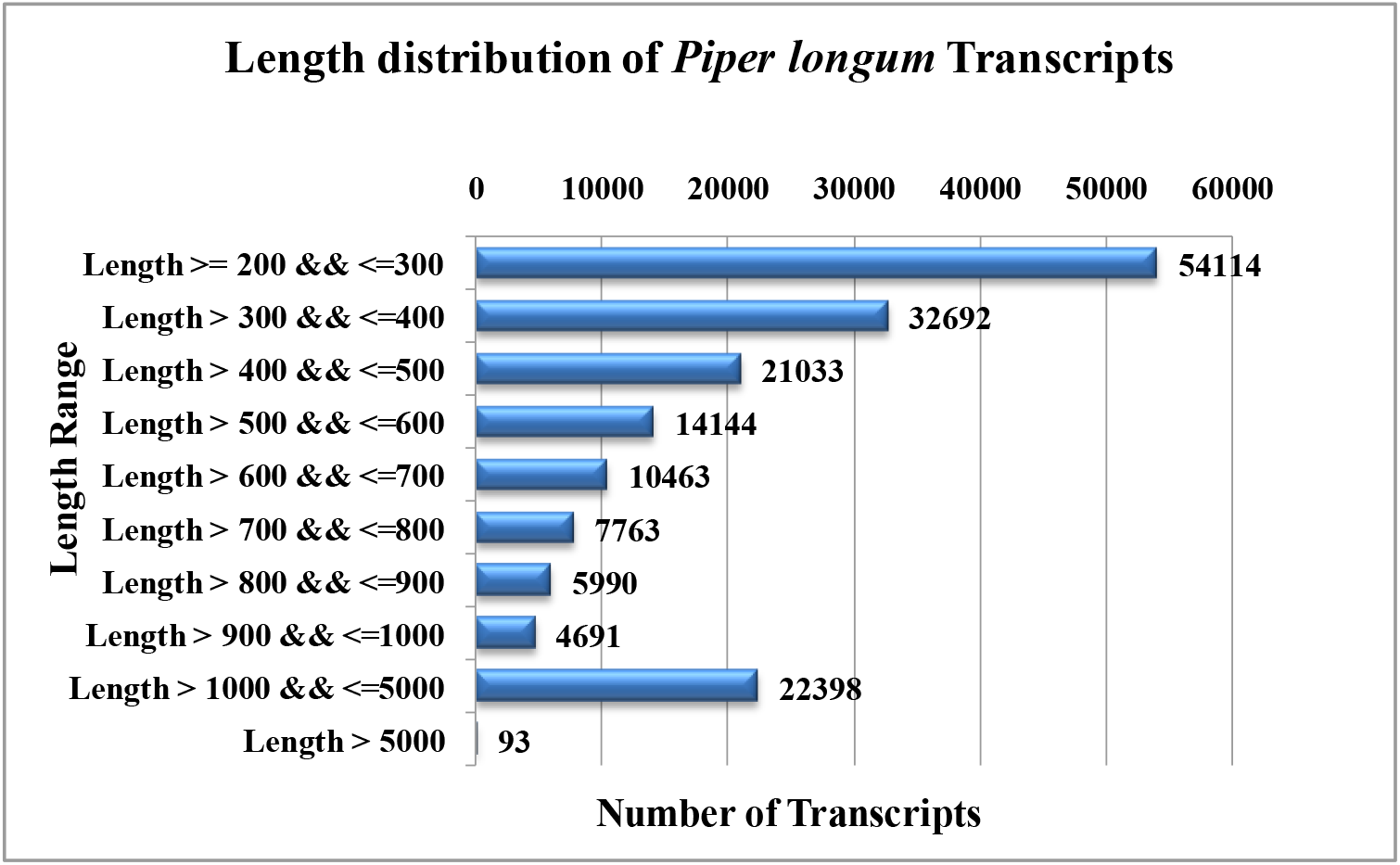
Length distribution of *Piper longum* transcripts.

### Feature of the combined assembly

To identify gene families of the samples of *P. longum*, the CDS obtained from assembly of each transcriptome were tagged, pooled and assembled again using Illumina TruSeq RNA sequencer. It has been considered that the super assembly of all primary assemblies is generated using different programs which provided better results in terms of consistency and size of transcripts. This analysis was carried out to get larger transcripts and to study digital differential expression of different tissues. CDS were predicted from the 1,73,381 assembled transcript sequences using Transdecoder (rel16JAN2014) at default parameters which resulted in identification of 58,773 CDS (Protein coding sequence) and total size of CDS was 3,87,34,722. The maximum and N50 length of CDS was 12990 and 747 bases. The size distribution of assembled CDS is displayed in Figure 2.

**Figure 2:**
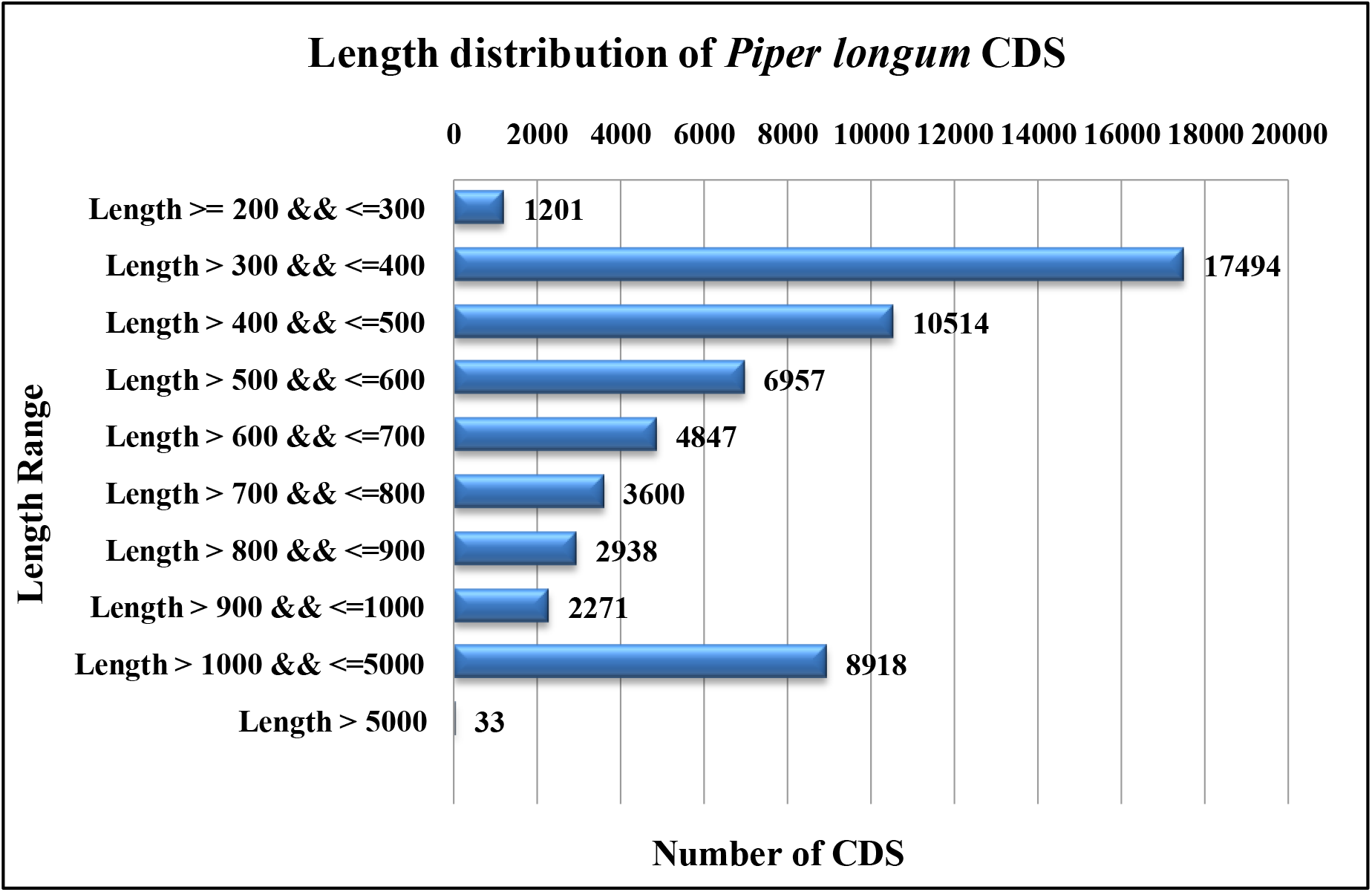
Length distribution of CDS of *Piper longum* sample.

### Functional Annotation of Predicted CDSs

Functional annotation of the CDS generated from the combined assembly of three different tissues of *P. longum* was carried out using Standalone BLASTx 2.2.30+. The annotation was performed on the predicted CDS of *P. longum* sample by aligning the CDS to non-redundant protein database of NCBI using BLASTx with e-value less than 1e-5. BLASTx analysis statistics of the predicted CDS with BLAST hits was 50277 and without BLAST hits was 8496 out of total 58773 CDS sequences (Table 2). Out of these different databases, top-BLAST hit was observed in *Nelumbo nucifera*, about 11934 CDS, followed by *Vitis vinifera* 3816 CDS and *Elaeis guineensis* 3496 CDS (Figure. 3).

**Table 2:**
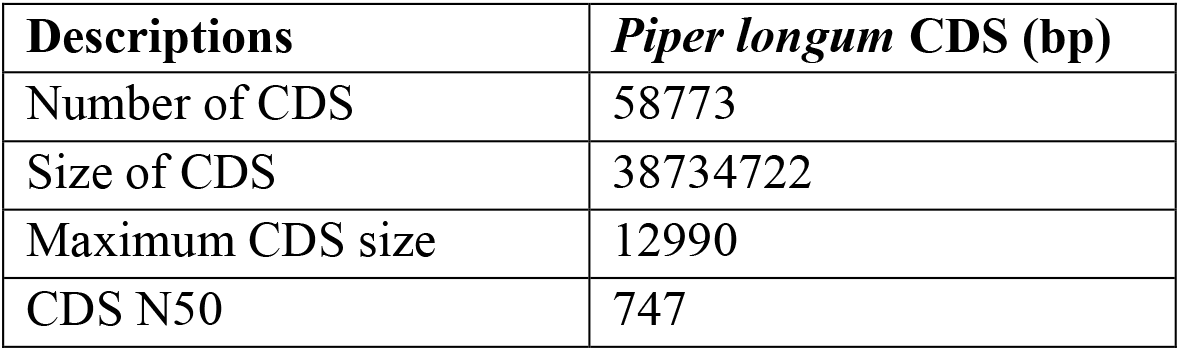
Statistics of predicted CDS of *Piper longum* sample.

**Figure 3:**
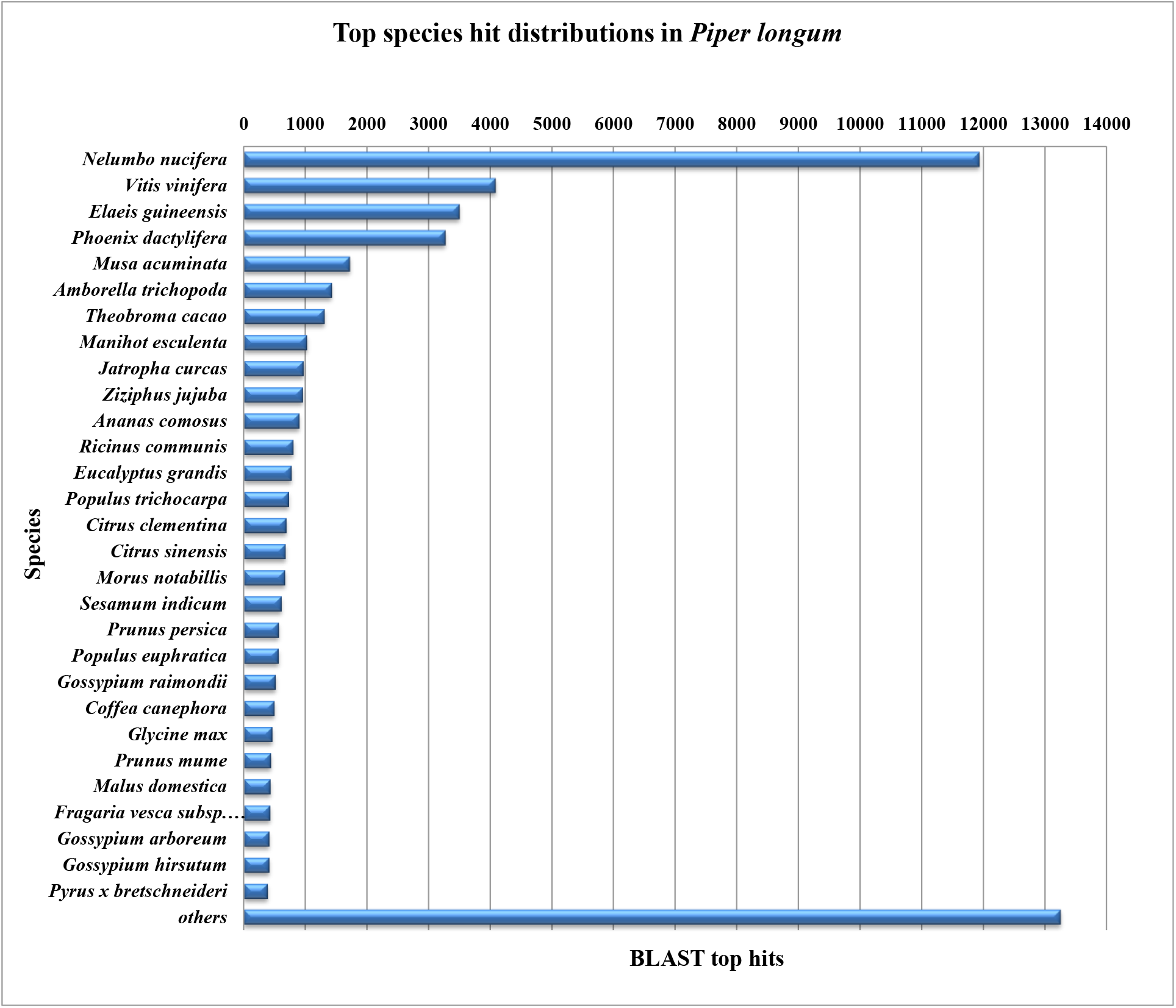
Top species hit distributions of *Piper longum* CDS.

### Gene annotation and functional classification

GO assignments were used to classify the functions of the predicted CDS. The GO mapping also provided ontology of defined terms representing gene product properties which were grouped into three main domains: Biological Process (BP), Molecular Function (MF) and Cellular Component (CC). BLASTx result accession IDs were searched directly in the gene product of GO database. As the international standardized gene functional classification system, GO and COG classifications were conducted for transcriptome data annotation. Among all of the categories, metabolic processes (24.6%) and cellular processes (21.9%) in the biological processes, membrane (13.8%) and cell (13.6%) in the cellular component, and binding (23.7%) and catalytic activity (22.4%) in the molecular function represented the major subcategories (Figure 4). In *P. longum*, 18508 CDS in biological process, 14429 CDS in Cellular component and 21103 CDS in Molecular function were obtained (Table 3).

**Figure 4:**
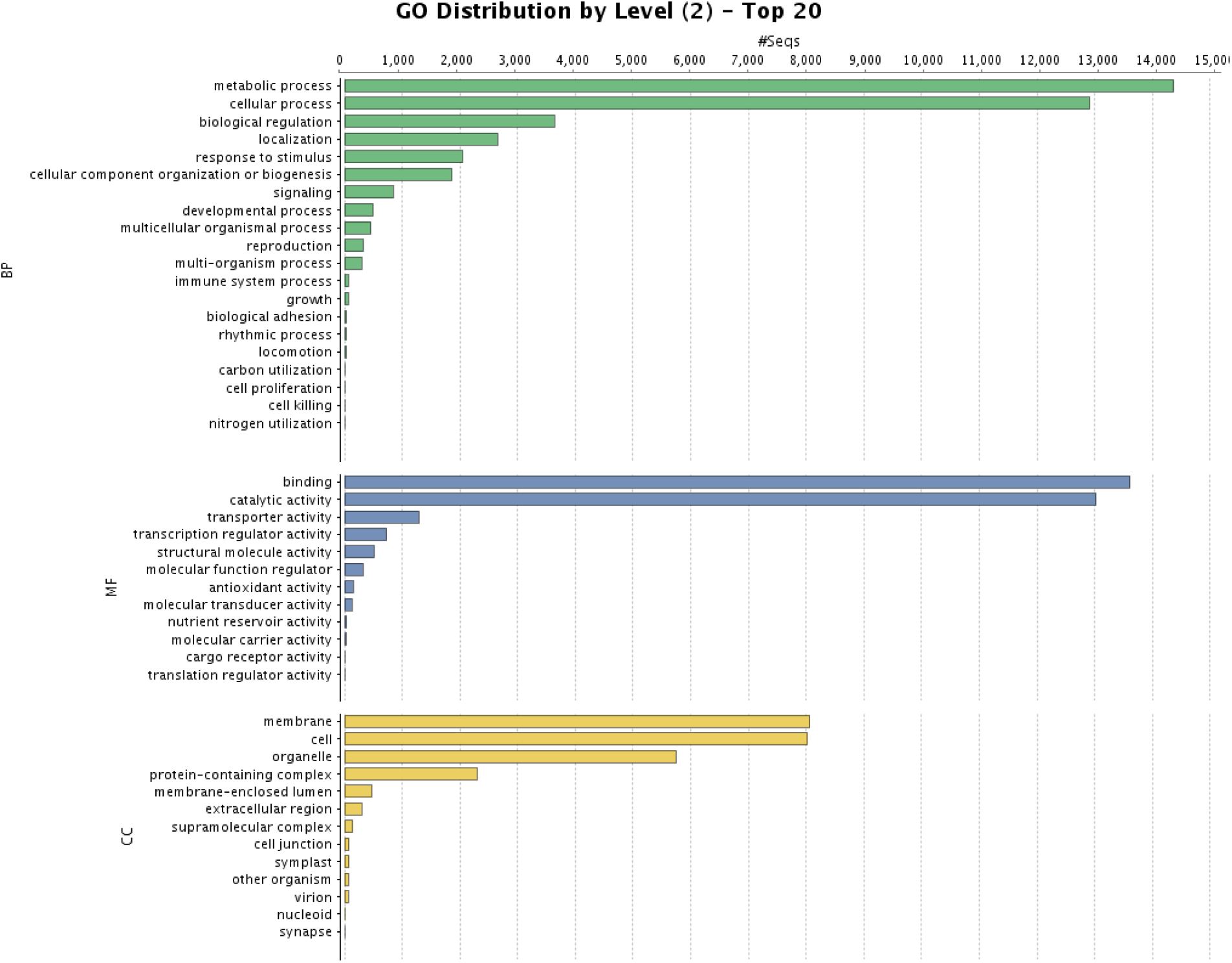
Gene ontology term (Biological process, Cellular Component and Molecular Function) distribution in *Piper longum* CDS.

**Table 3:**
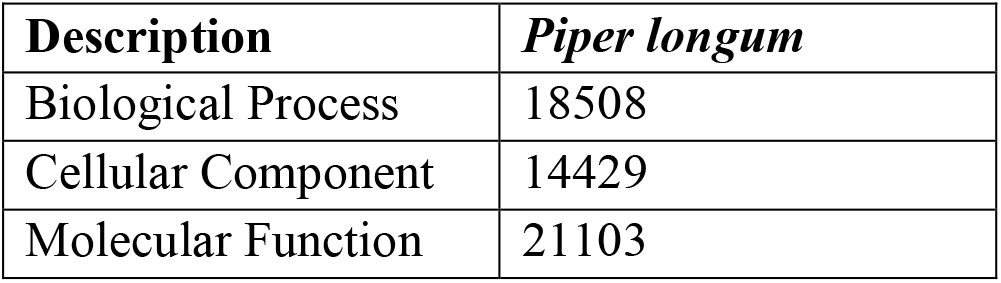
GO distribution statistics of the annotated CDS of *Piper longum*.

### Functional characterization and scanning of piperine-related gene using KEGG

KEGG pathway was used to identify the functional profiles of genes. The pathway analysis was carried out for all the samples using KEGG automatic annotation server. All the CDS were compared against the KEGG database using BLASTx with threshold bit-score value of 60 (default). The mapped CDS represented metabolic pathways of carbohydrates, lipids, nucleotides, amino acids, secondary metabolites, terpenoids, polyketides etc. The mapped CDS also represented the genes involved in genetic information processing, environmental information processing and cellular processes. A total of 2097 CDS (3.5%) were mapped into metabolism category followed by other pathways for *P. longum* (Figure 5). Among the CDS count 113 (0.2%), 290 (0.5%) and 103 (0.2%) were involved in biosynthesis of other secondary metabolites, amino acid metabolism and metabolism of terpenoids and polyketides, respectively. This data provided a valuable resource that might be used to study specific processes and pathways in the development of *P. longum*. Few genes were identified as piperidine pathway such as Lysine biosynthesis and Amino Acid biosynthesis related genes. In this pathway Tropane, piperidine and pyridine alkaloid biosynthesis [PATH: ko00960] represented 14 CDS.

**Figure 5:**
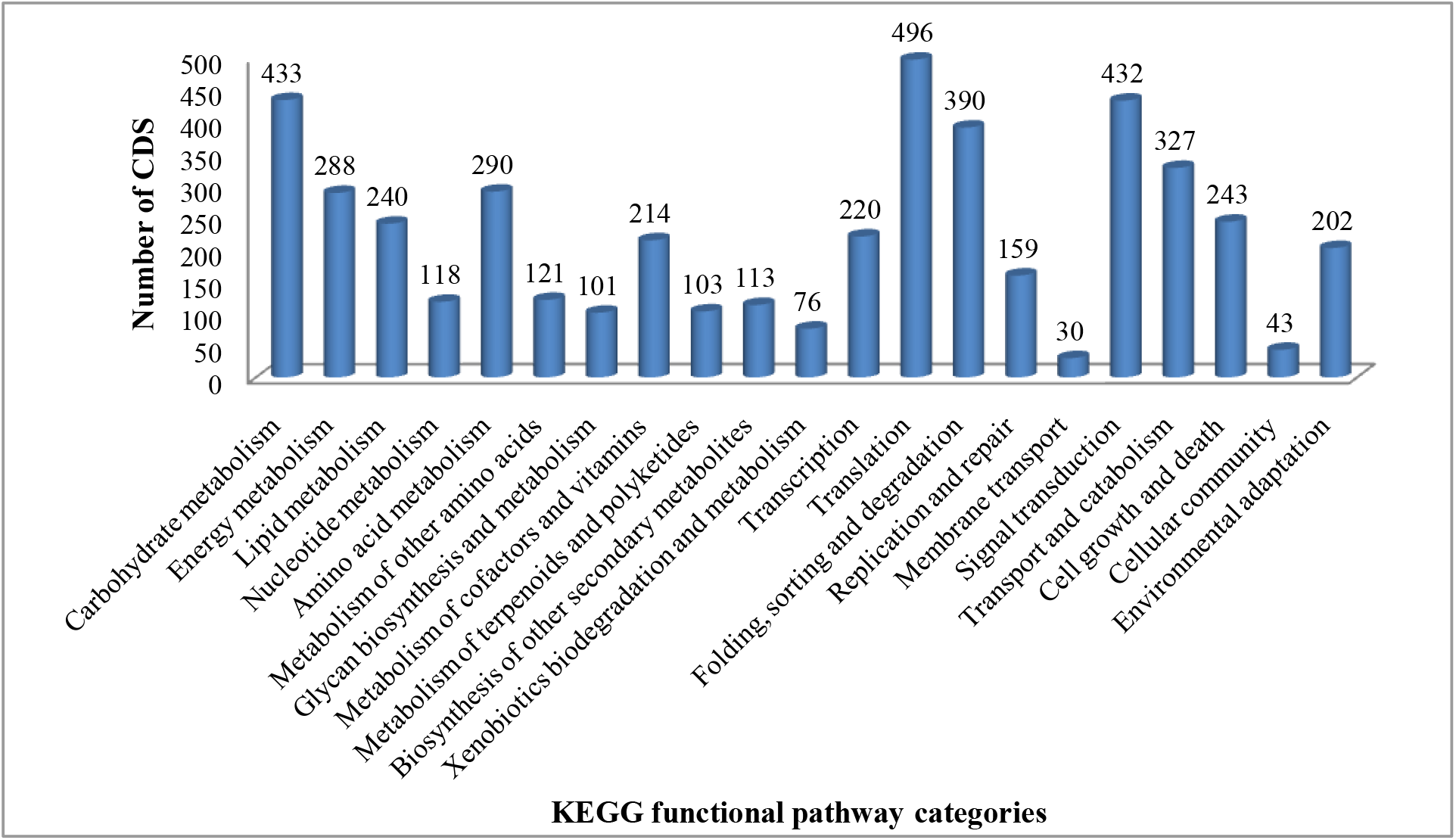
KEGG functional pathway of *Piper longum*.

### Differential gene expression analysis, primer designing and validation

Based on the common hits accession of functionally annotated CDS in samples, a total of 58773 CDS between leaf, root and spike were predicted. Differential gene expression analysis were divided into three different catagories: leaf vs root, leaf vs spike and root vs spike.

#### (a) Differential gene expression analysis between *P. longum* leaf (Control) and *P. longum* roots (Treated)

Differential gene expression analysis was carried out using commonly occurring NR blast hit accessions in the two samples. Within the set of 30296 commonly differentially expressed genes, a total of 581 genes were identified as significantly differentially up-regulated expressed and 1307 genes as significantly differentially down-regulated based on conditions p-value<0.05 (Table 4).

**Table 4:**
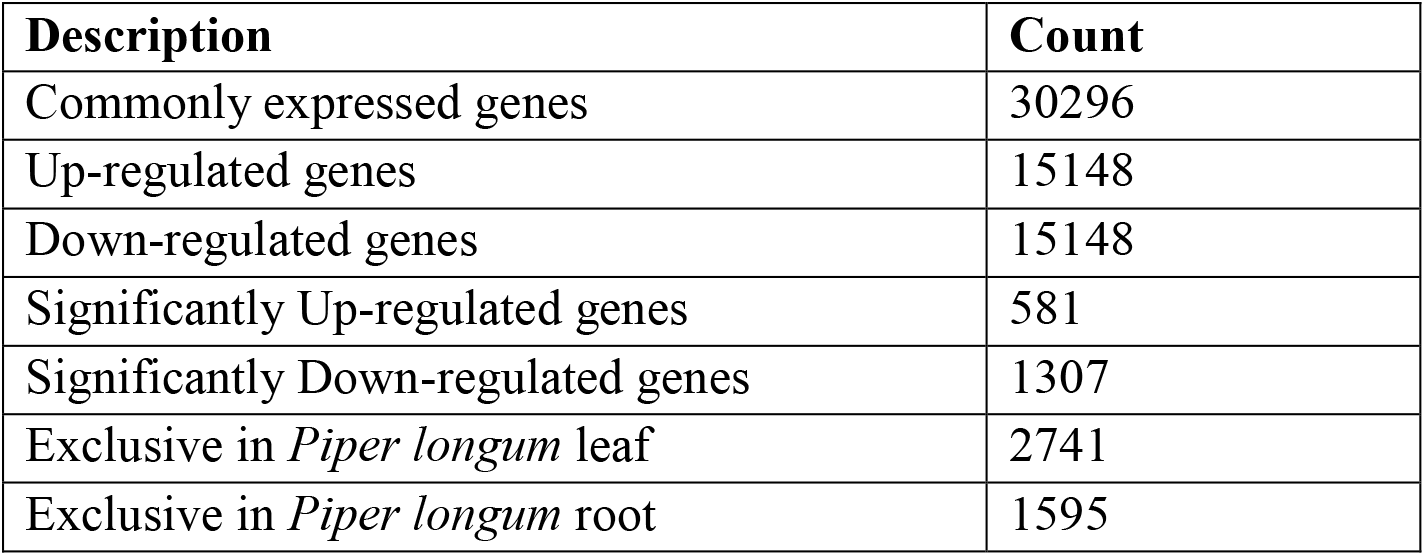
Distribution of genes expressed between *P. longum* leaf and root.

#### (b) Differential gene expression analysis between *P. longum* leaf (Control) and *P. longum* Spike (Treated)

Within the set of 32216 commonly differentially expressed genes, a total of 625 genes were identified as significantly differentially up-regulated expressed and 1246 genes as significantly differentially down-regulated based on conditions p-value<0.05 (Table 5).

**Table 5:** Distribution of genes expressed between *P. longum* leaf and spike.

| Description | Count |
| --- | --- |
| Commonly expressed genes | 32216 |
| Up-regulated genes | 16108 |
| Down-regulated genes | 16108 |
| Significantly Up-regulated genes | 625 |
| Significantly Down-regulated genes | 1246 |
| Exclusive in *Piper longum* leaf | 821 |
| Exclusive in *Piper longum* root | 1033 |

#### (c) Differential Gene Expression Analysis between *P. longum* roots (Control) and *P. longum* Spike (Treated)

Within the set of 30225 commonly differentially expressed genes, a total of 993 genes were identified as significantly differentially up-regulated expressed and 878 genes as significantly differentially down-regulated based on conditions p-value<0.05 (Table 6).

**Table 6:**
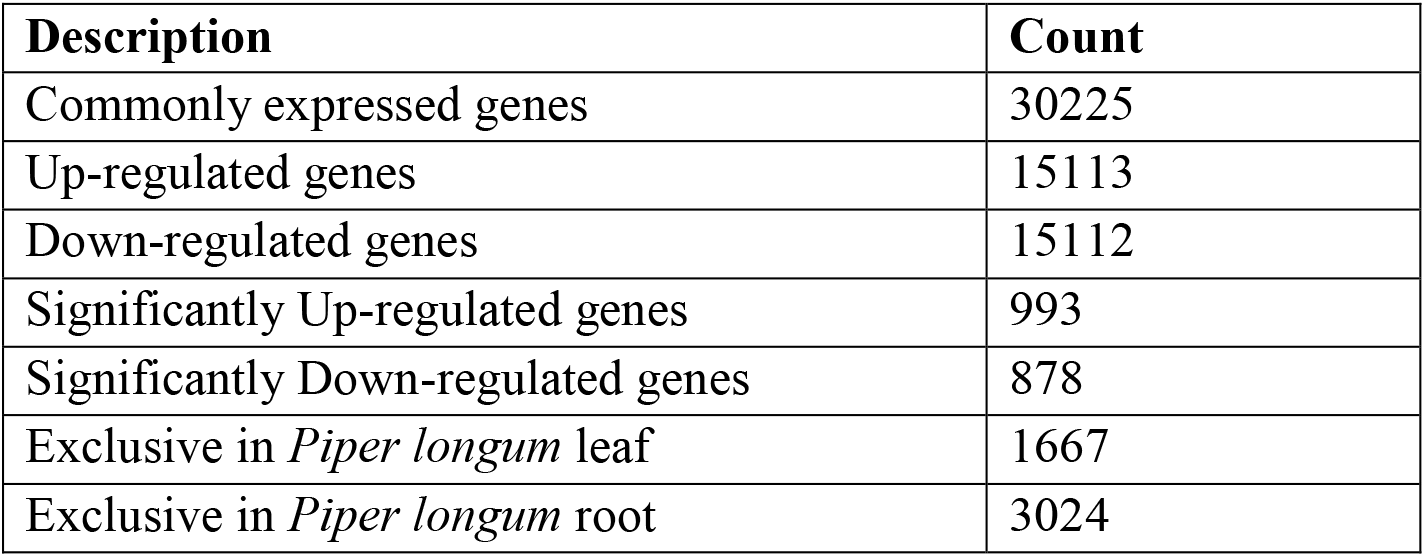
Distribution of genes expressed between *P. longum* root and spike.

### Detection of SSR markers

For detection of SSRs all the transcript contigs were searched with Perl script MISA (Microsatellite Searching Tool). A total of 8041 SSRs were found in assembled transcripts of *P. longum* (Table 7). Out of 8041 SSRs, 2619 SSRs were filtered to satisfy the 150 bp flanking criteria for *P. longum*. In these SSRs, the trinucleotide repeat motifs were the most abundant, accounting for 4506 SSRs (56.04%), followed by 3209 (39.91%) dinucleotide repeat motifs, 216 (2.69%) tetranucleotide repeat motifs, 79 (0.98%) pentanucleotide repeat motifs and 31 (0.38%) hexanucleotide repeat motifs (Table 8).

**Table 7:** SSRs identification statistics in *Piper longum*.

| Description | *P. longum* |
| --- | --- |
| Total number of sequences examined | 173381 |
| Total size of examined sequences (bp) | 99612836 |
| Total number of identified SSRs | 8041 |
| Number of SSR containing sequences | 7139 |
| Number of sequences containing more than 1 SSR | 737 |
| Number of SSRs present in compound formation | 570 |

**Table 8:**
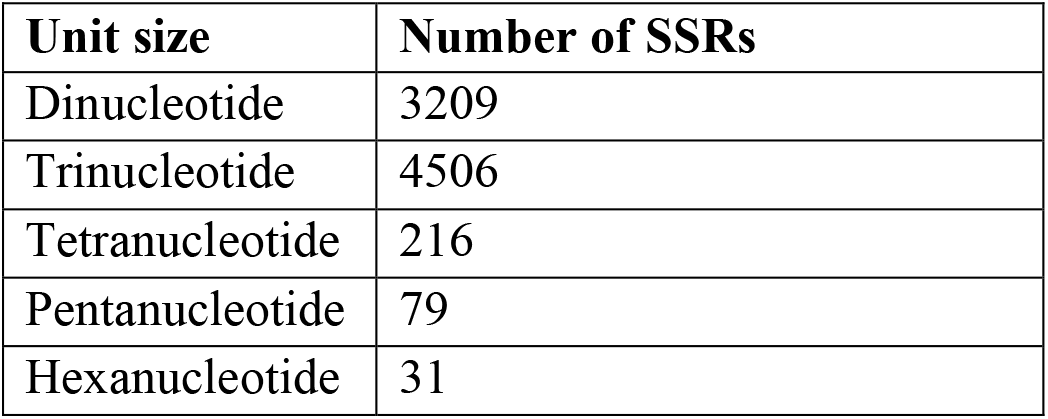
Distribution to different repeat type classes.

| Unit size | Number of SSRs |
| --- | --- |
| Dinucleotide | 3209 |
| Trinucleotide | 4506 |
| Tetranucleotide | 216 |
| Pentanucleotide | 79 |
| Hexanucleotide | 31 |

### Transcription factors (TFs)

Transcription factor encoding transcripts were analyzed by CDS comparison to known transcription factor gene families. In total, 21235 transcription factor genes, distributed in at least 65 families, were identified. In *P. longum* most abundant factors identified were bHLH, NAC and MYB_related which contain 2264, 1508 and 1370 CDS respectively. During the annotation analysis of the transcriptome data of *P. longum*, we have identified several TFs belonging to different families (Figure 6).

**Figure 6:**
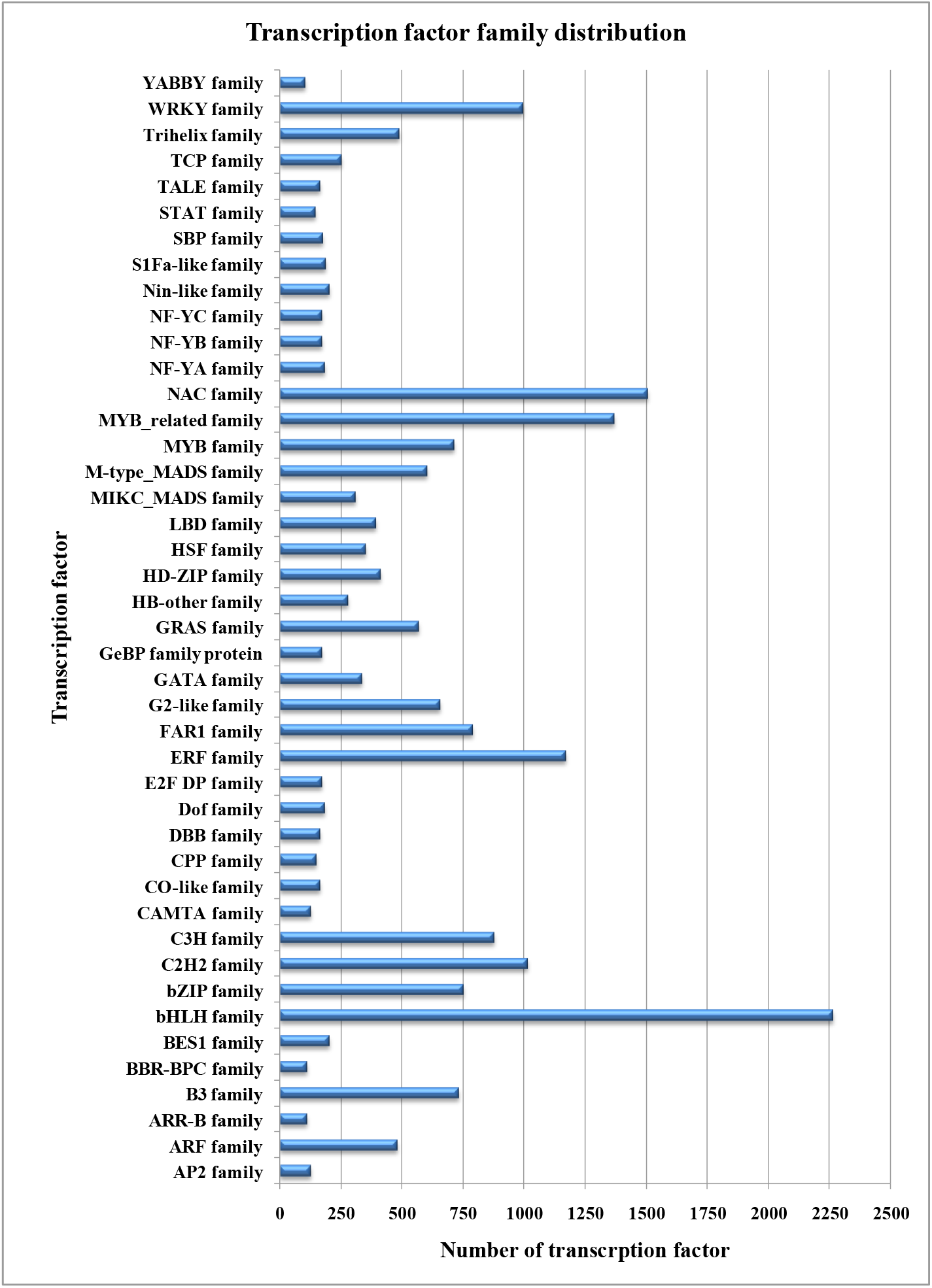
Distribution of different transcription factors in CDS of *Piper longum*.

### Identification of candidate genes involved in the piperine biosynthesis pathway

A total of 4730 CDS counts were assigned to 377 different pathways in KEGG database. Biosynthesis of secondary metabolites were categoried into 28 sections including phenylpropanoid biosynthesis (61 CDS), flavonoid biosynthesis (17 CDS), Tropane, piperidine and pyridine alkaloid biosynthesis (14 CDS), Isoquinoline alkaloid biosynthesis (11 CDS), Stilbenoid, diarylheptanoid and gingerol biosynthesis (9 CDS), Streptomycin biosynthesis (8 CDS) and biosynthesis of other secondary metabolites (Figure 8). Higher proportion of CDS belonged to the pathway of Translation, Signal transduction and Carbohydrate metabolism. Majority of CDS were classified into the metabolism, and the number of CDS related to different secondary metabolisms. To explore the regulatory mechanisms for the accumulation of piperine in *P. longum*, the expression profile of genes involved in piperine biosynthesis were analyzed (Figure 7). A total 14 expressed CDS encoding Tropane, piperidine and pyridine alkaloid biosynthesis enzymes were identified in *P. longum* (Table 9).

**Figure 7:**
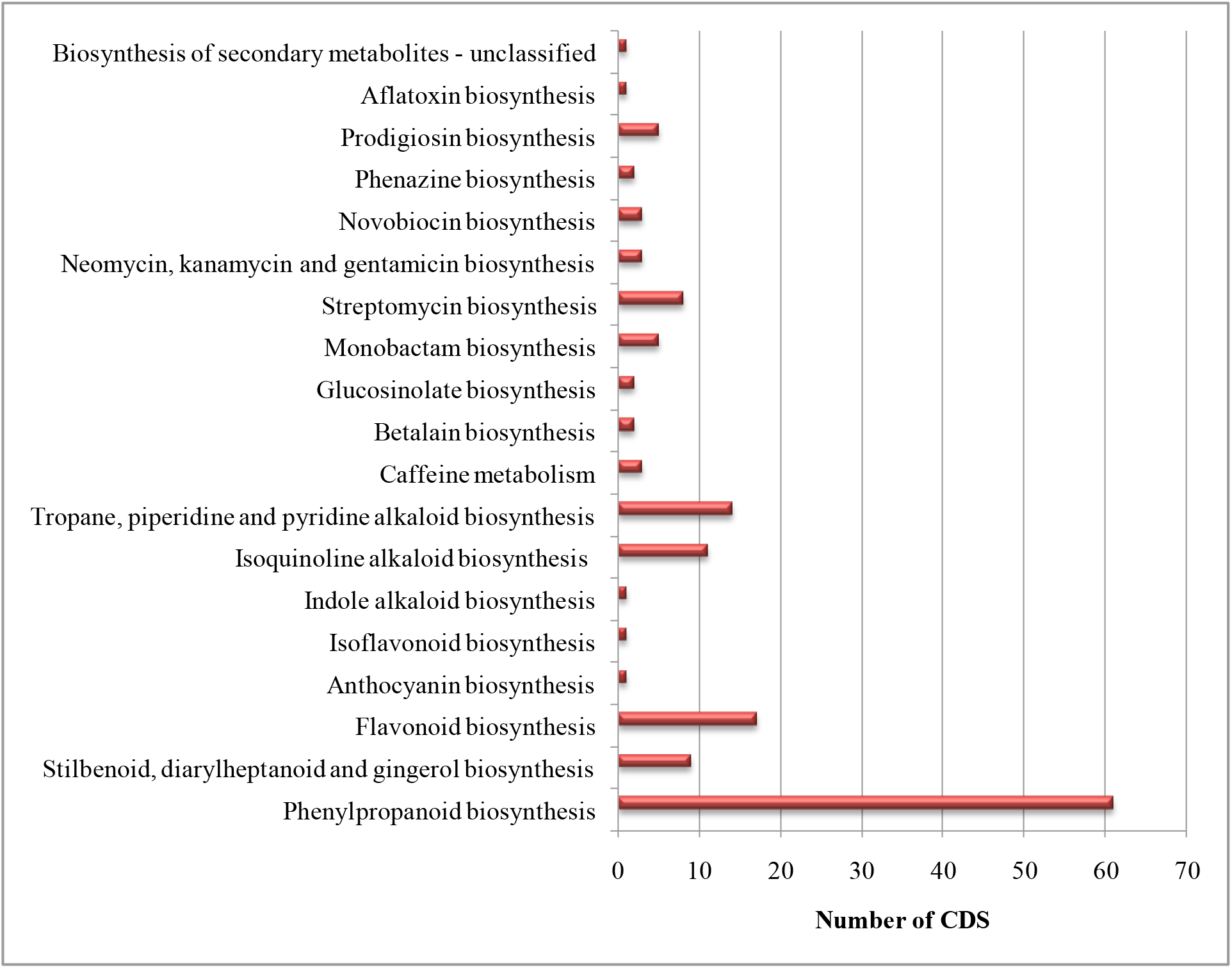
Classification based on categories of secondary metabolite biosynthesis.

**Figure 8:**
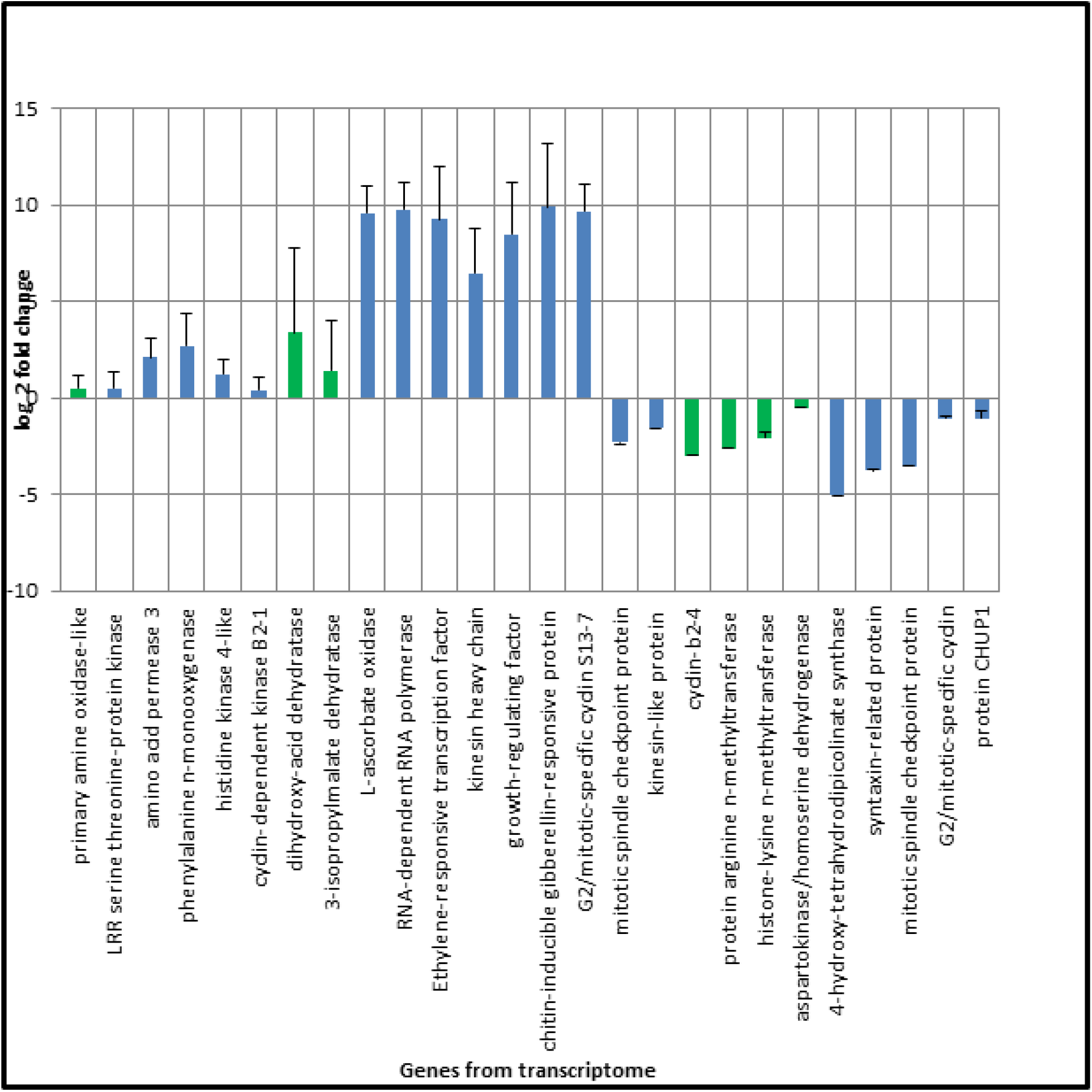
shows log2 fold change of selected differentially Expressed gene.

**Table 9:** CDS involved in Tropane, piperidine and pyridine alkaloid biosynthesis [PATH: ko00960].

| **00960 Tropane, piperidine and pyridine alkaloid biosynthesis [PATH: ko00960]** |  |  |  |  |
| --- | --- | --- | --- | --- |
| **S. No.** | **CDS IDs** | **KO IDs** | **EC IDs** | **Biosynthetic pathway** |
| 1 | cds_18211_Transcript_23293 | K00276<br>AOC3,<br>AOC2, tynA | primary-amine<br>oxidase<br>[EC:1.4.3.21] | Pyridine biosynthesis<br>pathway |
| 2 | cds_20102_Transcript_25925 | K00276<br>AOC3,<br>AOC2, tynA | primary-amine<br>oxidase<br>[EC:1.4.3.21] | Piperidine<br>biosynthetic pathway |
| 3 | cds_30149_Transcript_40912 | K00276<br>AOC3,<br>AOC2, tynA | primary-amine<br>oxidase<br>[EC:1.4.3.21] | Pyridine biosynthesis<br>pathway |
| 4 | cds_36462_Transcript_51168 | K00276<br>AOC3,<br>AOC2, tynA | primary-amine<br>oxidase<br>[EC:1.4.3.21] | Piperidine<br>biosynthetic pathway |
| 5 | cds_16196_Transcript_2043 | K08081<br>TR1 | tropinone reductase<br>I [EC:1.1.1.206] | Tropane biosynthetic<br>pathway |
| 6 | cds_56318_Transcript_925 | K08081<br>TR1 | tropinone reductase<br>I [EC:1.1.1.206] | Tropane biosynthetic<br>pathway |
| 7 | cds_56356_Transcript_926 | K08081<br>TR1 | tropinone reductase<br>I [EC:1.1.1.206] | Tropane biosynthetic<br>pathway |
| 8 | cds_16021_Transcript_2020 | K14454<br>GOT1 | aspartate<br>aminotransferase,<br>cytoplasmic<br>[EC:2.6.1.1] | Tropane biosynthetic<br>pathway |
| 9 | cds_39099_Transcript_55714 | K14454<br>GOT1 | aspartate<br>aminotransferase,<br>cytoplasmic<br>[EC:2.6.1.1] | Tropane biosynthetic<br>pathway |
| 10 | cds_42796_Transcript_6228 | K15849<br>PAT, AAT | bifunctional<br>aspartate<br>aminotransferase<br>and<br>glutamate/aspartate-<br>prephenate<br>aminotransferase<br>[EC:2.6.1.1<br>2.6.1.78 2.6.1.79] | Tropane biosynthetic<br>pathway |
| 11 | cds_42801_Transcript_6229 | K15849<br>PAT, AAT | bifunctional<br>aspartate<br>aminotransferase<br>and<br>glutamate/aspartate-<br>prephenate<br>aminotransferase<br>[EC:2.6.1.1<br>2.6.1.78 2.6.1.79] | Tropane biosynthetic<br>pathway |
| 12 | cds_20398_Transcript_26340 | K00815<br>TAT | tyrosine<br>aminotransferase<br>[EC:2.6.1.5] | Tropane biosynthetic<br>pathway |
| 13 | cds_22551_Transcript_29424 | K00815<br>TAT | tyrosine<br>aminotransferase<br>[EC:2.6.1.5] | Tropane biosynthetic<br>pathway |
| 14 | cds_13069_Transcript_16650 | K00817<br>hisC | histidinol-<br>phosphate<br>aminotransferase<br>[EC:2.6.1.9] | Tropane biosynthetic<br>pathway |

### Validation of Expression of Differentially Expressed genes-

To validate differential expression, 27 genes were selected (Table 10, 11) to confirm the expression level by Quantitative Real Time PCR (qRT-PCR) m. Validation of expression of selected genes through qRT-PCR suggests that differentially expressed genes identified through digital gene expression analysis might be differentially expressed between samples (Figure 8).

**Table 10:** List of genes differentially expressed in leaf and spike selected for validation and its differential expression in terms of log 2 fold.

| S. No. | Name of Genes | Function of Genes in Leaf | Function of Genes in Spike | log2fold |
| --- | --- | --- | --- | --- |
| 1 | GAPDH | Internal control gene | Internal control gene | 0 |
| 2 | 4-hydroxy-tetrahydrodipicolinate synthase | Absent | Lysine biosynthesis | -5.058892212 |
| 3 | Amino acid permease 3 | Absent | Lysine biosynthesis | 2.114903455 |
| 4 | Aspartokinase/homoserine dehydrogenase | Aspartate pathway | Absent | -0.505489973 |
| 5 | LRR receptor-like serine threonine-protein kinase | Absent | Aspartate pathway | 0.504469951 |
| 6 | 3-isopropylmalate dehydratase | Absent | biosynthesis of leucine | 1.423867816 |
| 7 | Phenylalanine n-monooxygenase | Absent | Valine, leucine and isoleucine pathway | 2.715696386 |
| 8 | Histidine kinase 4-like | Absent | biosynthesis of leucine | 1.23091656 |
| 9 | Dihydroxy-acid dehydratase | Absent | Valine, leucine and isoleucine pathway | 3.408627507 |
| 10 | Primary amine oxidase-like | Piperidine Pathway related enzyme | Absent | 0.479662132 |
| 11 | Protein arginine n-methyltransferase | Regulating flowering time | Absent | -2.610880587 |
| 12 | Histone-lysine n-methyltransferase | Regulating flowering time | Absent | -2.040028228 |
| 13 | L-ascorbate oxidase | Absent | Ascorbate metabolism | 9.574864201 |
| 14 | RNA-dependent RNA polymerase | Absent | Replication of RNA from an RNA template. | 9.766782064 |
| 15 | Ethylene-responsive transcription factor | Absent | Stress signal transduction pathways. | 9.234066916 |
| 16 | Kinesin heavy chain | Absent | Calmodulin-binding protein | 6.475303198 |
| 17 | Growth-regulating factor | Absent | Regulating flowering time | 8.460050976 |
| 18 | Chitin-inducible gibberellin-responsive protein | Absent | Regulating flowering time | 9.902196359 |
| 19 | Cyclin-dependent kinase B2-1 | Absent | Ascorbate metabolism | 0.389143966 |
| 20 | Kinesin-like protein | Replication of RNA from an RNA template. | Absent | -1.568212769 |
| 21 | Cyclin-b2-4-like | Stress signal transduction pathways. | Absent | -2.973046787 |
| 22 | G2/mitotic-specific cyclin S13-7 | Calmodulin-binding protein | Absent | 9.662897235 |
| 23 | Mitotic spindle checkpoint protein | Microtubule generation, bipolarity establishment, length control and chromosome alignment | Absent | -3.512551913 |
| 24 | Syntaxin-related protein | Syntaxin localized at the plasma membrane and cell signaling | Absent | -3.749175868 |
| 25 | Mitotic spindle checkpoint protein | Microtubule generation, bipolarity establishment, length control and chromosome alignment | Absent | -3.512551913 |
| 26 | G2/mitotic-specific cyclin | Accumulate during G2 and destroyed at mitosis | Absent | -1.067365167 |
| 27 | Protein CHUP1 | For actin polymerization and chloroplast relocation movement | Absent | -1.058641104 |

**Table 11:**
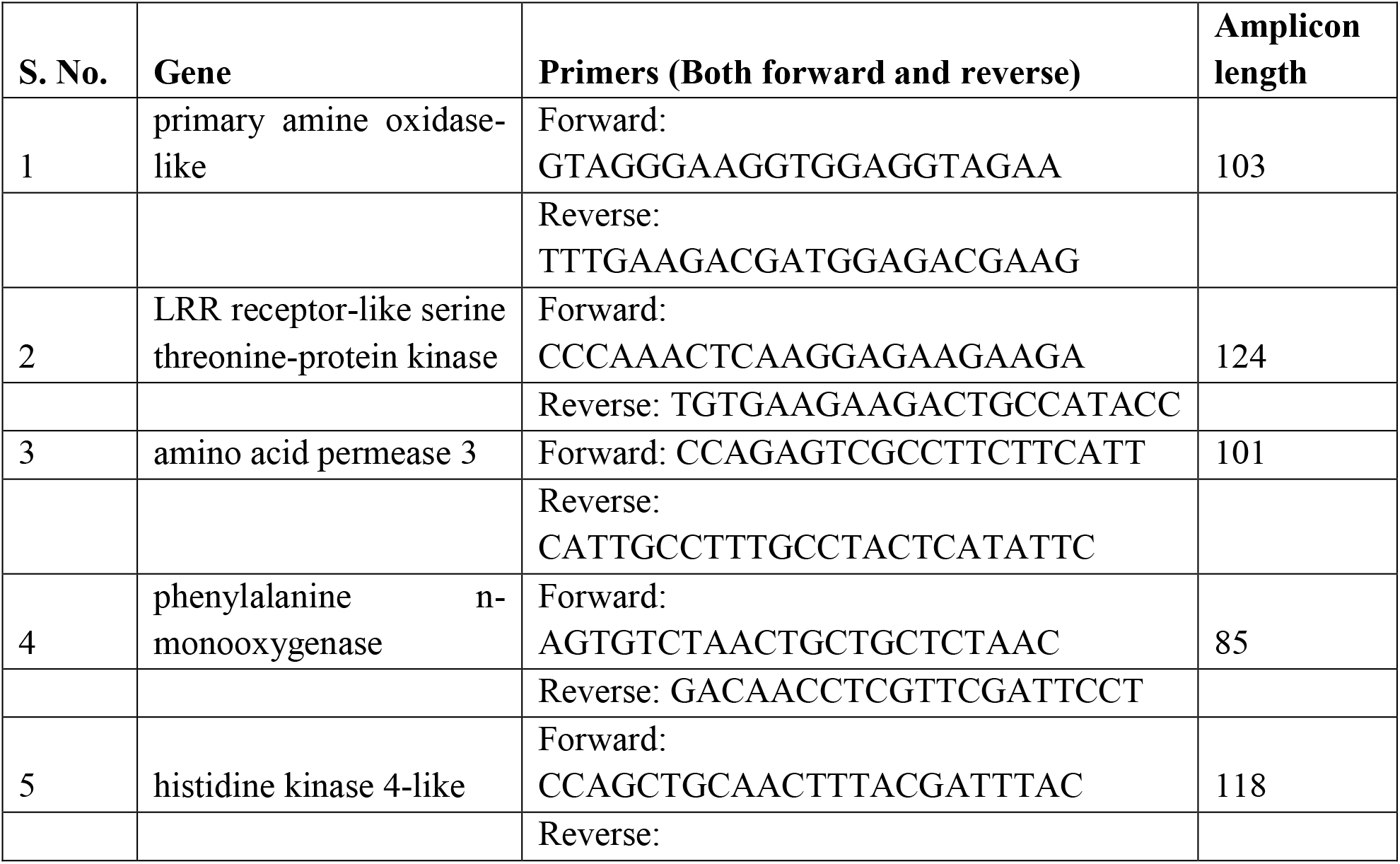
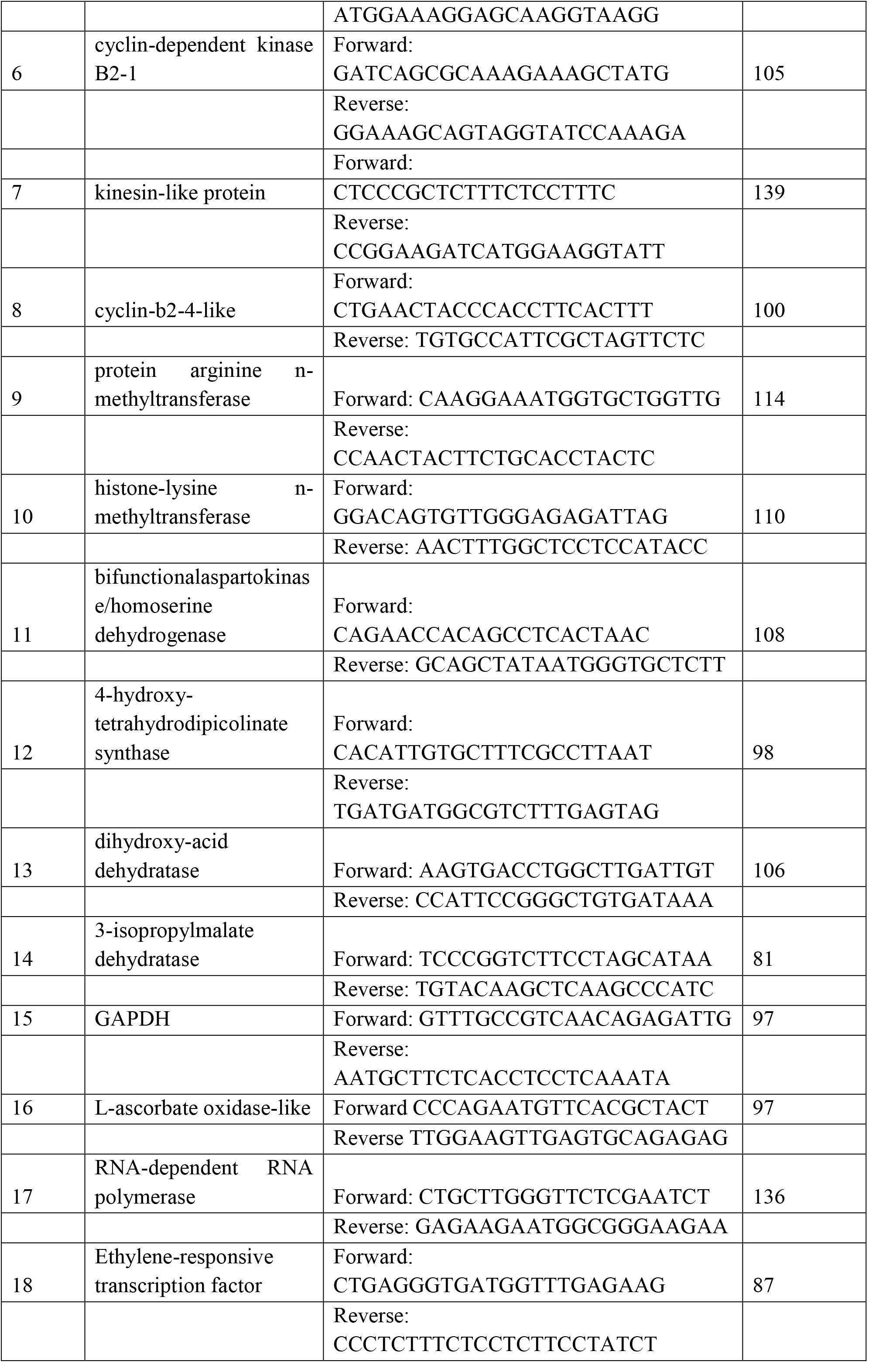
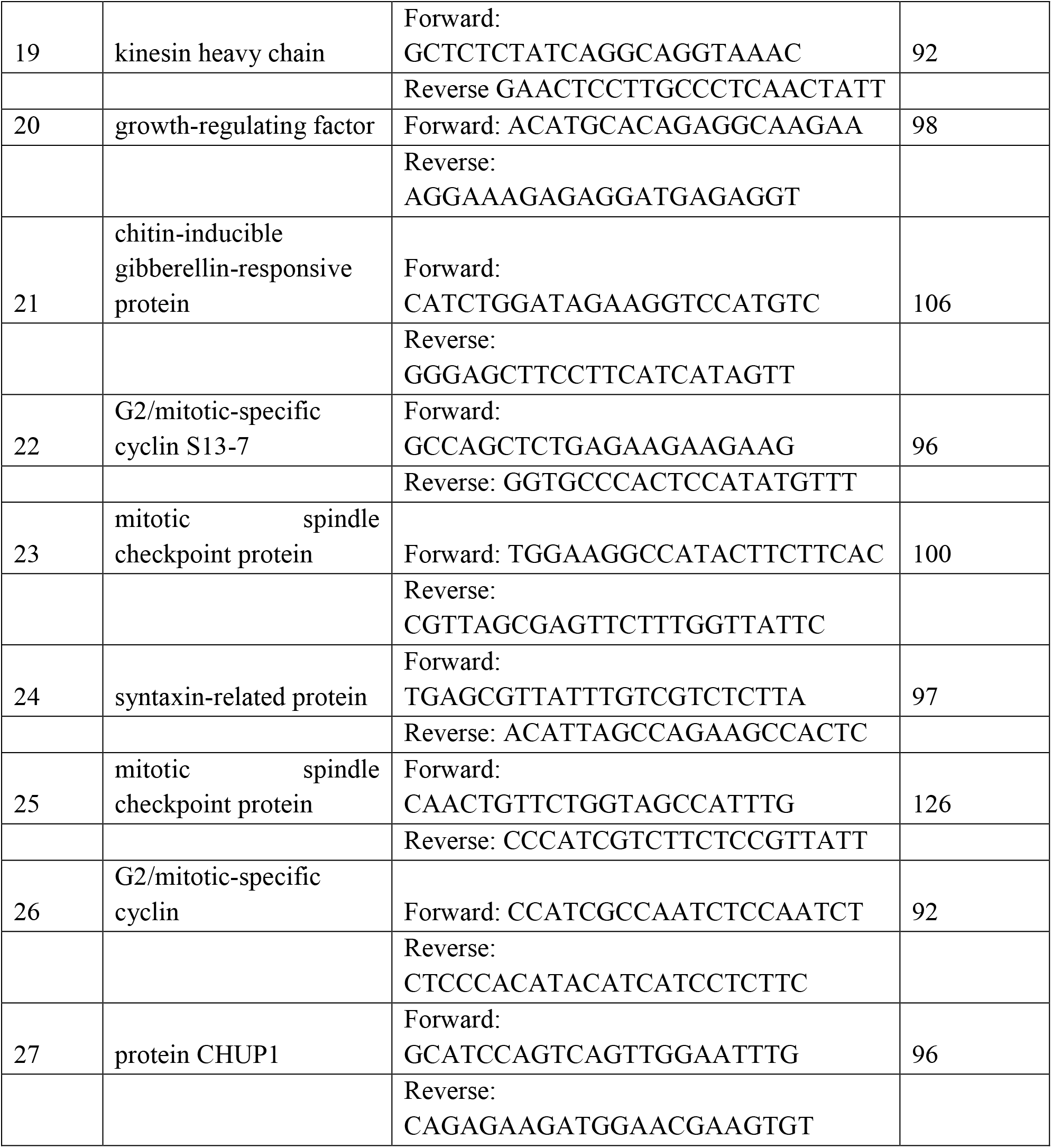
List of selected primers for validation of Expression of Differentially Expressed gene.

### Development piperine biosynthesis-related gene-

Piperoyl CoA and piperidine are the two precursor of piperine using acyltransferase. However, piperine is related with three main groups:

Group A: Gene associated with phenylpropanoid pathway (map00940). In this pathway cinnamoyl-CoA is developed for piperoyl CoA biosynthesis.

Group B: Genes related with L-lysine metabolism (map01064). In this pathway lysine is converted into piperidine using series of reaction.

Group C: Genes allied with acyltransferase, which perform catalytic activity in between piperoylcoenzyme A and piperidine.

The spike of plant shows the importance of arogenate dehydratase (ADT), amino transferase (PPA-AT), p-coumarate 3-hydroxylase (C3H), cinnamate 4-hydroxylase (C4H/CYP7) HCT, caffeic acid-3-O-methyltransferase (CAOMT), p-coumarate-CoA-ligase (4CL). All these genes are concerned in the reaction of phenylpropanoids to piperoyl CoA. The concern transcripts associated with different enzymes and compounds are shown in the Table 12. Finally, the piperine synthesis and their related genes are observed expansion of multiple gene families.

**Table 12:** Transcripts associated with different enzymes and compounds.

| S. No. | Name | No. of Transcript |
| --- | --- | --- |
| 1 | Shikimate | 38 |
| 2 | Shikimate kinase (2.7.1.71) | 4 |
| 3 | Shikimate 3-phosphate | - |
| 4 | 3-Phosphoshikimate 1-carboxyvinyl transferase (2.5.1.19) | 3 |
| 5 | 5-0-(1-Carboxy vinyl)-3-phosphoshikimate | 3 |
| 6 | Chorismate synthase (4.2.3.5) | 7 |
| 7 | Chorismate | 1 |
| 8 | Chorismate mutase (5.4.99.5) | 4 |
| 9 | Prephenate | 24 |
| 10 | Aromatic-amino-acid transaminase (2.6.1.57, 2.6.1.1, | 10 |
|  | 2.6.1.78, 2.1.1.79) |  |
| 11 | L-Arogenate | - |
| 12 | Prephenate dehydratase (4.2.1.51, 4.2.1.91, 4.2.1.51) | 3 |
| 13 | L-Phenylalanine | 16 |
| 14 | phenylalanine ammonialyase (4.3.1.24, 4.3.1.25) (PAL) | 18 |
| 15 | trans-cinnamate | - |
| 16 | trans-cinnamate 4-mono-oxygenase (1.14.14.91) (C4H) | 5 |
| 17 | 4-coumarate or p-coumaric acid | - |
| 18 | p-coumarate 3-hydroxylase (1.14.13.-) (C3H) | 2 |
| 19 | Caffeate / Caffeic acid | - |
| 20 | caffeic acid 3-O-methyltransferase (2.1.1.68) (CAOMT) | 6 |
| 21 | Ferulate or Ferulic acid | - |
| 22 | 4-coumarate CoA ligase (6.2.1.12) (4CL) | 12 |
| 23 | Feruloyl-coA | 4 |
| 24 | cinnamoyl CoA reductase (1.2.1.44) | 1 |
| 25 | Coniferyl aldehyde | - |
| 26 | 4-coumarate coA ligase (6.2.1.12) (4CL) | 12 |
| 27 | Shikimate-O-hydroxy cinnamoyl transferase (2.3.1.133) (HCT) | 3 |
| 28 | Caffeoyl coA | - |
| 29 | 5-O-caffeoyl shikimic acid | 15 |
| 30 | Chaffeoyl quinic acid | - |
| 31 | 5-O-(4-coumaroyl)- D-quinic acid-3'- monooxygenase (1.14.14.96) | 3 |
| 32 | p-Coumaroyl quinic acid | - |
| 33 | 4-coumaroyl CoA | - |

## Discussion

In nature 12,000 alkaloids are known and many more are yet to be discovered (Facchini *et al*., 2012). Transcriptome databases of specific plant species generate numbers of biosynthetic genes encoding enzymes with valuable catalytic activities and variants with similar functions, but different biochemical features. As compared to complete genome sequencing, transcriptome sequencing is a good substitute for rapid and efficient access to the expressed genes and characterizing phenotypes (Wang *et al*., 2010). There is a benefit of *de novo* assembly and gene models could be obtained that otherwise may not be assembled by any other method as the reads may not be mapped back to the current genomic scaffolds due to improper scaffold assembly. A differential expression of gene analysis for different tissues was investigated in the present study with Digital Gene Expression (DGE) profiling technology (Strickler *et al*., 2012) to understand the functional genomics for *P. longum*. Previous studies showed that de novo transcriptome assembly using high quality Illumina RNA-Seq reads were more useful for genome assembly. However, if the gene transcripts are fragmented and gene annotation is incomplete, de novo assembly can help in the reading of gene map back to the assembly and also in gene model representation. If a genome assembly is present, it can be useful to unite *de novo* when the databases are available (Huang *et al*., 2016).

For Next Generation Sequencing (NGS) with the help of RNA-Seq data, transcriptome assembly is the first important step. However, in non-model plants NGS is a challenge, especially due to the high level of ploidy, large gene size, enormous quantity of data, large proportion of repeat sequences and low genome complexities produced by transcriptome assembly (Wang *et al*., 2010). Aim of our study was to provide important and useful information to the database which is helpful to the researchers accomplishing transcriptome analysis in non-model plants. Gene transcripts show better expression characteristics in plants and also provide metabolic engineering alternative for optimization of synthetic biosystems designed to produce valuable plant metabolites (Xiao *et al*., 2013).

The Next Generation Sequencing study in non-model plants as *P. longum* will lead to identification of pathways and gene related to production of natural products such as piperine and would also help to discover additional pathways. Transcriptome analysis is cost effective approach of sequence determination and improves the efficiency and speed of gene discovery (Jose and Sharma, 1985; Vasavirama and Upender, 2014). *P. longum* is an important plant used for medicinal purposes but transcriptome and genetic information are not available in NCBI database. Leaves, roots and spikes of this plant are used for medicinal purpose. Despite having immense medicinal value, there is very little information available regarding biosynthesis of piperine from this plant (Vasavirama and Upender, 2014) and lack the complete genetic data (Paterson *et al*., 2006; Jiao *et al*., 2011; Sheila *et al*., 2012). In GenBank database, genome of black pepper (*P. nigrum*) is represented by only 184 sequences (Jaramillo *et al*., 2001; Menezes *et al*., 2009; Sen *et al*., 2010; Joy *et al*., 2007).

In the comparative characterizations of the leaf, root and spike transcriptome of *P. longum* valuable resources were generated for new gene discovery and development of SSR markers for further study. To the best of our knowledge, there is no other report on Illumina paired-end sequencing technology for *P. longum* leaf, root and spike transcriptome de novo sequencing and assembly without reference genome. The results demonstrated that Illumina paired end sequencing can be used as a fast and efficient approach to gene discovery and molecular marker development for non-model organism, especially those with larger genome (Wang *et al*., 2010; Martin and Wang, 2011; Xiao *et al*., 2013; Xu *et al*., 2015; Hagel *et al*., 2015; Jain, 2011). This information represents a major trancriptomic level resource for *P. longum* and will be useful for comparative transcriptomic studies in *P. longum*. These analyses would provide genomic information to understand patterns of adaptive variation across the genome and to identify the genetic basis of adaptive traits (Parchman *et al*., 2010).

In the present analysis, a total of 99 million high quality reads were generated from leaves, roots and spikes of *P. longum* using Illumina TruSeq RNA. From these reads, 1,73,381 transcripts were sequenced and analyzed generating 8,041 SSRs. It gives a new fingerprinting for taxonomic contrasts, species mapping for this species. These analyzed data would help to describe the mechanisms and genes involved in alkaloid biosynthesis in non-model plants. Transcriptomic studies on medicinal plants as *Cassia angustifolia* (Reddy *et al*., 2015), *Panax vietnamensis* (Zhang *et al*., 2015), *Phyllanthus amarus (*Bose and Chattopadhyay, 2016) and *Chelidonium majus* (Pourmazaheri *et al*., 2019) has revealed important information regarding biosynthetic pathways of the active principles present in these plants. In these studies, only few transcripts were annotated by GO distribution as limited database is available for non-model plant sequences resulting in low hits in gene identification. SSRs are more informative and reliable markers in organism analysis (Idrees and Irshad, 2014; Madhumati, 2014; García *et al*., 2018; Ahmad *et al*., 2018).

The functional annotation of the dataset of CDS revealed synthesis of Lysine in *P. longum*. In spike, the activity of amino acid especially Lysine is very high as compared to leaf and root due to presence of piperine. Several differentially expressed transcripts were identified and analyzed in the three plant tissues of *P. longum*. Comparative transcriptome study of *P. longum* identified exclusive gene in leaf (2741 CDS) and root (1595 CDS) by the investigation of leaf and root tissue sample. Same analysis was found in leaf (821 CDS) and spike (1033 CDS) by the comparative analysis of both these tissues. Again a differential gene expression analysis identified exclusive genes in root (3024 CDS) and spike (1667 CDS).

Gene Ontology (GO) unites genes and gene product attributes across all the species (The Gene Ontology project, 2008) and also provideds information about the role of metabolic alkaloids. GO analysis identified 28 transcripts involved in the secondary metabolite pathway specially related to the piperine production including Lysine biosynthesis. In future, these transcripts will be important resources for genetic manipulation of *P. longum*. The metabolic pathway of piperine are not known and little information is available on biosynthesis of tropane, piperidine and pyridine alkaloid biosynthesis in plants (Sui *et al*., 2011, Liu *et al*., 2015; Zhang *et al*., 2015; Liao *et al*., 2016; Mazumdar and Chattopadhyay, 2016; Liu *et al*., 2017; Shou-ming *et al*., 2017; Li *et al*., 2018; Lei *et al*., 2018).

Piperine is a lysine-derived alkaloid and in the first step Lysine through decarboxylation produces cadaverine (Bunsupa *et al*., 2014). Cadaverine is converted into piperidine (Wink and Hartmann, 1979, Wink, 1987, Okwute *et al*., 2013). Some of CDS related to enzymes involved in these steps also showed different expression (Gordo *et al*., 2012 and Joy *et al*., 2013). The number of Illumina reads in each CDS is directly proportional to the abundance of specific cDNA in the library. This study helped us to study differential expression of genes related to pathway and their specific role in tissue-specific piperine biosynthesis and molecular mechanism of leaf, root and spike tissue. TFs control gene expression in response to various internal and external signals by suppressing and enhancing downstream genes (Reddy *et al*., 2015). These factor families known to manage secondary metabolism play an important role in regulation of piperine biosynthesis. TFs family plays critical roles in interactions with other molecules (Takatsuji, 1998; De *et al*., 1999).

A large number of SSR markers were identified from the transcriptome data and it is an important molecular tool to be used in gene mapping, genetic diversity and molecular breeding. SSR markers also provide useful reference data for further research. Transcriptional factors regulate gene expression of external and internal signals for activating or suppressing downstream genes (De Folter *et al*., 2005). Transcripts were found to be putative transcription factor encoding regions not belonging to any particular transcription factor family. These families were known to regulate secondary metabolism and play important role in piperine biosynthesis. C3H transcription factors belong to zinc finger motif family that plays a major role in interactions with other molecules (Takatsuji, 1998; De *et al*., 1999). Functional characterization of the candidate genes will not only help the biochemical mechanism of compound, but also provide molecular and biochemical target for biotechnological interventions.

In this study, comparative study of piperine biosynthesis related with tropane, piperidine and pyridine alkaloid biosynthesis pathway were addressed, and 15 functional genes were found (Reddy *et al*., 2015, Ishihara *et al*., 2015, Sun *et al*., 2017 and Lei *et al*., 2018). In the first attempt, more genes were significantly down-regulated than up-regulated as compared with leaf and root. In the second attempt, again more genes were down-regulated than up-regulated as compared with leaf and spike. In the last attempt, more genes were up-regulated than down-regulated as compared with root and spike. Pathway based analysis for the leaves and spike of *P. longum* is helpful to further understand the function and gene interactions. Lysine are the most important and basic structure for the piperine biosynthetic pathway (Szoke et al., 2013, Hu et al., 2015, Chopra, et al., 2016). In biosynthetic process of piperine, N-heterocycle piperidine and thioester piperoyl-CoA was transferring their group (Geissmann and Crout, 1969). The decarboxylation of L-lysine in the presence of pyridoxal phosphate (PLP) formed cadaverine. Then oxidative deamination via enzyme oxidative deamination was produced amino aldehyde and after cyclization provides imine, Δ1-piperideine. And the product were further reduced to piperidine and after the reaction with piperoyl-CoA final product piperine was created (Simon and Henry, 2013). Still this process has not been proven yet, thus many enzymatic analysis depend upon the biosynthesis of similar compounds like coumaroylagmatine (Bird and Smith, 1983, 1989) and feruloyltyramine (Negrel and Martin, 1984, Negrel and Jeandet, 1987) were carried out. But the reaction between piperoyl-coenzyme A and 2-piperidine that produces piperine was shown in the shoots of *P. nigrum* (Semler et al., 1897; Semler and Gross, 1988).

The pathway of piperine is starts with shikimate being converted to shikimate 3-phospthate by shikimate kinase (2.7.1.71), and formed 5-O-(1-carboxyvinyl)-3-phosphoshikimate with the help of 3-phosphoshikimate 1-carboxyvinyl transferase (2.5.1.19) (KEGG pathway) (Legrand et al., 2016). Then by using chorismate synthase (4.2.3.5), chorismate produced. L-phenylalanine is synthesized after two steps from chorismate (Figure 9). The phenylalanine being converted to cinnamic acid by phenylalanine ammonia lyase (PAL), and diverges into several branches at q-coumaroyl CoA (Dewick, 2002; Ehlting et al., 2006). The first enzyme in the phenylpropanoid pathway is phenylalanine ammonia-lyase (PAL). Three other enzymes of phenylpropanoid and lignin metabolism, were shown to increase, notably 4-coumarate: CoA ligase (4CL), caffeic acid/5-hydroxyconiferaldehyde O-methyltransferase (COMT) and cinnamyl alcohol dehydrogenase (CAD) (Hoffmann et al., 2003). Thus, 4 enzymes involved in the phenylpropanoid and lignin pathways are PAL, p-hydroxycinnamoyl-CoA: quinate shikimate p-hydroxycinnamoyltransferase (HCT), cinnamoyl-CoA reductase (CCR) and CAD (Andersen et al., 2008; Peng et al., 2008).

**Figure 9:**
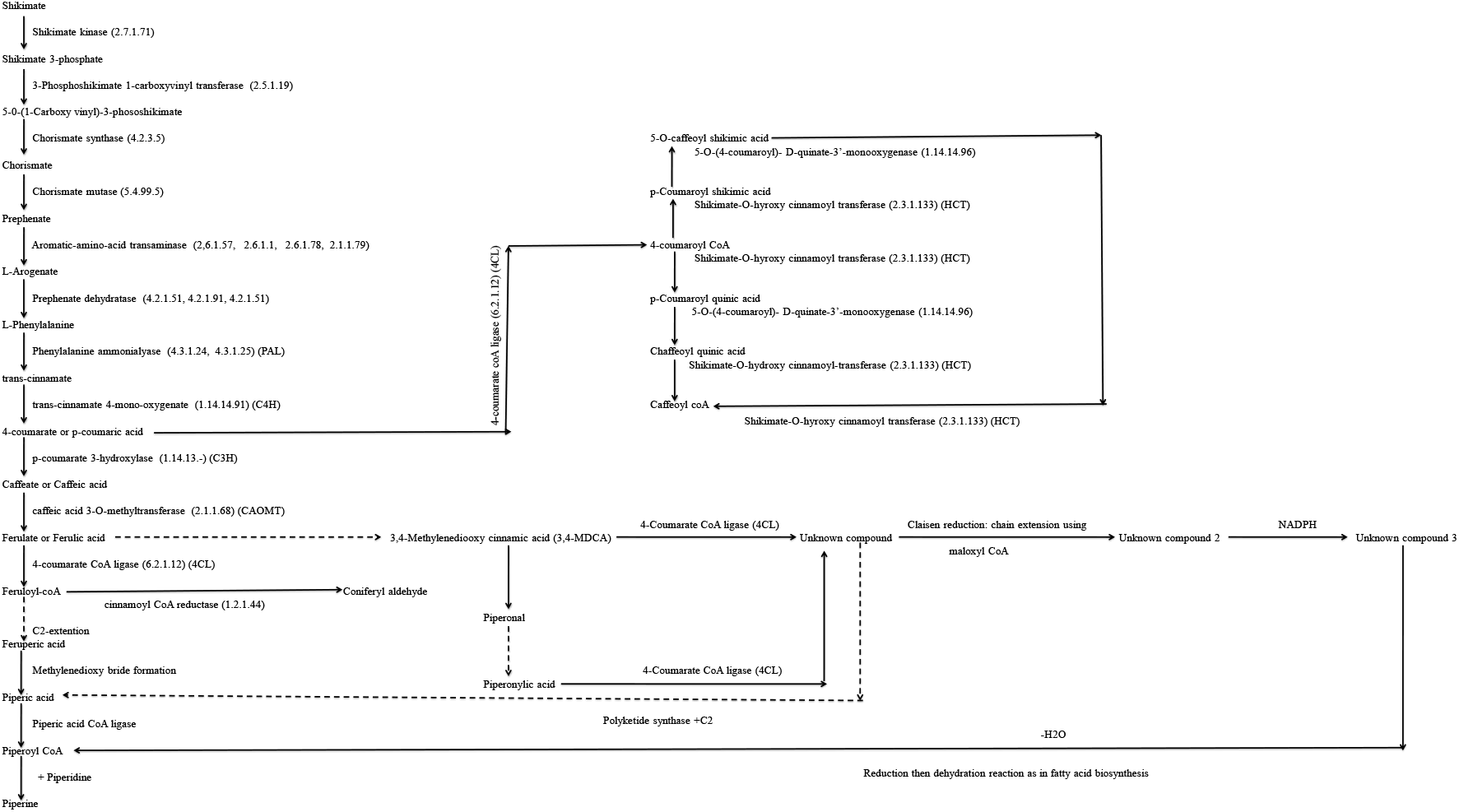
Piperine biosynthetic pathway.

Piperine consists of 3,4-methylenedioxyphenyl piperidine connected by a C5 amide bridge. 3,4-Methylenedioxyphenyl and part of the bridging chain is known to bifurcate from the phenylpropanoid pathway (Richet et al., 2012). The biosynthesis of piperine in that 3,4-methylenedioxycinnamoyl-CoA, derived through phenylpropanoid pathway, would undergo one C2 chain extension using malonyl-CoA to arrive at piperoyl-CoA (Schnabel et al., 2020). The presence of (E,E)-piperoyl-CoA: piperidine N piperoyltransferase, catalyzing the ultimate reaction in the piperine pathway to form piperine in the shoots of the plant (Guzman, 2014). The above-mentioned putative pathway proposed for CoA thioester formation of phenylpropenoate(s) at the certain stage of phenylpropanoid pathway to activate the carboxyl group of the acid(s) (Chen et al., 2013). This type of activation is commonly carried out by 4-coumarate: CoA ligase (4CL) (Jin et al., 2020). Members in 4CL family can be classified into three subfamilies according to their catalytic functions and amino acid sequences.

Class I: 4CLs are typically involved in lignin biosynthesis, whereas

Class II: 4CLs participate in plant’s response to environmental stress and in the biosynthesis of specialized metabolites.

Class III: Coumarate:CoA ligase-like (CLL) clade was recently described

In piperine, the aromatic part of the molecules is linked by an amide bond piperidine. The piperidine heterocycle is derived from L-lysine, whereas the aromatic part of piperine is likely to be derived from phenylpropanoid metabolism. Piperoyl-CoA is required for the piperine synthase activity. Although the acetyl or malonyl CoA-based elongation of feruloyl-CoA appears to be a possible mechanism for the C2-extension of the cinnamic acid-derived precursor resulting in piperoyl-CoA formation. Based on computational analysis, we now report the piperic acid is accepted as a substrate. PipCoA ligase might be required for the accumulation of piperine during early fruit ripening and may provide the piperoyl-CoA ester for the subsequent piperine transferase reaction, the key to piperine.

In an article the spike specific expression related genes contains lysine decarboxylase (LDC), glycosyltransferases (GFTs), cytochrome P450s (CYP); shikimate hydroxycinnamoyl transferases (HCT) in the phenylpropanoid pathway. The amino acid pathways, acyltransferases (ATs) and phenylpropanoid are ubiquitous in plant secondary metabolism (Hu et al., 2019). In case of Capsicum species, the capsaicinoids are derived from phenylpropanoid (Aza-González et al., 2011). Lysine-derived quinoline alkaloids are create by the synthesis of phenylpropanoid and lysine metabolism in Papaver somniferum, Macleaya cordata (Liu, X. et al., 2017), Nelumbo nucifera and Carica papaya (Bennett et al.,1997).

From the functional annotation of *P. longum* plant two genes were found related to the pyridoxal phosphate (PLP) viz., Cys Met pyridoxal phosphate-dependent enzyme and Pyridoxal phosphate (PLP)-dependent transferases superfamily isoform 1. Both are important genes and they may help in further study of piperine biosyhthesis.

The shape of inflorescence architecture is depended on the meristems produced in the inflorescence apex which defines the relative position of flowers. In *P. longum* plant the fruit is a spike form of inflorescence due to their floral transition. The development of inflorescence may be mostly expressed by the function and mutual combination of three genes: APETALA 1 (AP1), LEAFY (LFY) and TERMINAL FLOWER 1 (TFL1) (Shannon and Meeks-Wagner, 1993; Liljegren *et al*., 1999; Blazquez *et al*., 2006, Benlloch *et al*., 2015, Zhang *et al*., 2015, Cheng *et al*., 2018, Ma *et al*., 2018). These genes help to mentain the balance between the floral meristem and inflorescence (Alvarez *et al*., 1992, Ratcliffe *et al*., 1999; Blazquez *et al*., 2006). LFY gene is involved in the identification, regulation and development of floral meristem (Schultz and Haughn, 1991, Shannon and Meeks-Wagner, 1993). In our study, functional annotation of the transcripts revealed two genes related to LFY gene, one gene for LEAFY and again to gene function like TFL gene. The inflorescence of plant is a complex network that entails multiple pathways. Abundant genes actively participate in the process of inflorescence development, viz *STERILE APETALA* (*SAP*), *LEAFY* (*LFY*) and few floral homeotic genes, *AGAMOUS* (*AG*) and *APETALA2* (*AP2*). In C Class, gene AG, LFY is an up-regulated and in Class A, AP2 and SAP are down-regulated, which is involved in the development of reproductive organ (Krogan *et al*., 2012). These genes, we have found, are present in piper plant as well with four CDS in SAP, eight CDS in APETALA2 and seven CDS in AGAMOUS. Both WUSCHEL (WUS) and ULTRAPETALA 1 genes play important roles in flowering process such as activation of AG and ovule development (Lenhard et al., 2001). In the present study 21 annotations have been assigned to WUSHEL genes while three annotations have been assigned to ULTRAPETALA 1. AGL (MADS-box) transcripts are necessary for the development of reproductive organs (de Folter *et al*., 2005). In our study 19 genes have been annotated to AGL and 35 genes involved in LRR-RLK. GIGANTEA gene help in the end control of flowering, Circadian Clock and 27 annotations have been assigned to GIGANTEA in our study on *P. longum* (Peláez et al., 2019, Zhou et al., 2019). By using KEGG mapping of the best hits CDS, we have identified large number of CDS involved in different process viz. genetic information processing, metabolism, cellular processes, environmental information processing and organizational systems.

However, synthesis of piperine is extracted from decarboxylation and amine oxidation of lysine. And also they recommended as polymerisation of two tyrosines in benzylisoquinoline alkaloid biosynthesis. The derivative of phenylpropanoid and lysine metabolism is the two precursors which produce piperine after catalysed by acyltransferase. Thecombination of phenylpropanoid and lysine metabolism particularly decarboxylation and amine oxidation of lysine and acyl transformation reveal the feature of piperine synthesis.

Sequencing of long pepper has developed our consideration of the different piperine biosynthesis of Piper longum. Our analysis provides valuable impact that may create a base for the future research.

## Acknowledgments

Authors are thankful to Director and Head Department of Botany, Dayalbagh Educational Institute, Dayalbagh, Agra for providing infrastructure and support to carry out the research work. Mrinalini Prasad is grateful to Department of Science and Technology, New Delhi for research fellowship (DST-INSPIRE).

